# Human pericytes degrade α-synuclein aggregates in a strain-dependent manner

**DOI:** 10.1101/2022.06.08.495286

**Authors:** Birger Victor Dieriks, Blake Highet, Ania Alik, Tracy Bellande, Taylor J. Stevenson, Victoria Low, Thomas I-H Park, Jason Correia, Patrick Schweder, Richard L. M. Faull, Ronald Melki, Maurice A. Curtis, Mike Dragunow

**Affiliations:** Department of Anatomy and Medical Imaging, University of Auckland Private Bag 92019, Auckland, 1142, New Zealand; Centre for Brain Research, University of Auckland, Private Bag 92019, Auckland, 1142, New Zealand; Molecular Imaging Research Center, Francois Jacob Institute, Alternative Energies and Atomic Energy Commission, and Laboratory of Neurodegenerative Diseases, National Center for Scientific Research, Fontenay- Aux-Roses, France; Department of Pharmacology, University of Auckland, Private Bag 92019, Auckland, 1142, New Zealand; Auckland City Hospital, Park Road, Auckland, New Zealand

**Keywords:** pericytes, α-synuclein aggregates, strains, degradation

## Abstract

Parkinson’s disease (PD) is a progressive, neurodegenerative disorder characterised by the abnormal accumulation of α-synuclein (α-syn) aggregates. Central to disease progression is the gradual spread of pathological α-syn. α-syn aggregation is closely linked to progressive neuron loss. As such, clearance of α-syn aggregates may slow the progression of PD and lead to less severe symptoms. Evidence that non-neuronal cells play a role in PD and other synucleinopathies such as Lewy body dementia and multiple system atrophy are increasing. Our previous work has shown that pericytes — vascular mural cells that regulate the blood-brain barrier — contain α-syn aggregates in human PD brains. Here, we demonstrate that pericytes efficiently internalise fibrillar α-syn irrespective of being in a monoculture or mixed neuronal cell culture. Pericytes efficiently break down α-syn aggregates *in vitro*, with clear differences in the number of α-syn aggregates/cell and average aggregate size when comparing five pure α-syn strains (Fibrils, Ribbons, fibrils65, fibrils91 and fibrils110). Furthermore, pericytes derived from PD brains have a less uniform response than those derived from control brains. Our results highlight the vital role brain vasculature may play in reducing α-syn burden in PD.

## INTRODUCTION

Parkinson’s disease (PD) is a progressive, degenerative disorder of the brain that primarily affects the dopaminergic neurons in the substantia nigra and causes characteristic movement symptoms. Pathologically, PD is grouped with other synucleinopathies such as Lewy body dementia (LBD) and multiple system atrophy (MSA), which are all characterised by the abnormal accumulation of α-synuclein (α-syn) aggregates. Current data suggests that under normal physiological circumstances, monomeric cytosolic α-syn exists in a transient state until the protein interacts with membranes where it plays a role in synaptic transmission, vesicle endocytosis and membrane remodeling [1]. However, in people with PD, an event takes place that allows the formation of an initial α-syn oligomer or aggregate [2]. Once an initial α-syn oligomer is established, the oligomer functions as a seed to form larger and more extensive aggregates such as Lewy bodies and Lewy neurites. This process of aggregate formation has an important role in α-syn pathogenicity and progressive neurodegeneration in PD [2]. In most instances, cells can break down these aggregates through chaperone and proteasome machinery (reviewed in [3]). However, progressive α-syn aggregation may exceed the clearance capacity of affected cells. Aging further reduces these protective mechanisms, thereby enhancing aggregate accumulation and spread to neighbouring cells, where the process is repeated. Over the course of several years α-syn aggregates slowly spread into other brain regions, causing the gradual onset of characteristic PD symptoms [2,4].

This mode of action explains the gradual onset of disease, but it does not explain the variability in cell types affected and symptoms observed in patients with PD, LBD and MSA [5]. Several studies have identified noticeable differences in structural and phenotypic traits of fibrillar α-syn aggregates concerning seeding capacity and neurotoxicity. This led to the hypothesis that different α-syn aggregate 3D conformations or ‘strains’ may be partly responsible for the heterogeneous nature of PD and other synucleinopathies [6,7]. Several studies show that injection of pure α-syn strains or brain extracts isolated directly from human PD and other synucleinopathies into the rodent brain results in differential cortical propagation of α-syn pathology and reproduces the heterogeneity in pathology and symptoms observed in the original patients. The structure of injected strains remains unchanged throughout this process, emphasising the link between strain and patient symptomatology [7–10].

Neurons are the primary source of endogenous α-syn within the brain, and in PD, many develop large α-syn aggregates or Lewy bodies. However, non-neuronal cells have become increasingly linked to PD, with α-syn aggregates found in astrocytes and microglia [11,12]. Our recent quantification study of human PD olfactory bulbs shows that the number of non-neuronal cells (including astrocytes, microglia and pericytes) containing small aggregates is similar to the number of neurons with small aggregates [13]. Astrocytes and microglia are involved in phagocytic clearance of α-syn in a coordinated attempt to avoid propagation of PD pathology to neighbouring neurons [14–16]. However, nothing is known about the potential α-syn clearance role of pericytes in the brain.

Pericytes are contractile cells that regulate blood flow and permeability of the blood-brain barrier. Pericytes play an important role in neuroinflammation, acting as targets and early initiators of brain inflammation in response to systemic inflammation prior to astrocyte and microglia activation [17– 19]. PD causes loss of pericyte coverage of vessels and leakiness of the blood-brain barrier [20,21]. Furthermore, pericytes overexpressing α-syn in the cytoplasm can transfer α-syn through nanotubes that connect neighbouring cells [22]. Like microglia and astrocytes, pericytes could contribute a neuroprotective role in the clearance of α-syn aggregates, thereby reducing α-syn induced toxicity, as well as slowing down spread [14,15]. Therefore, we studied pericyte α-syn degradation by exposing cultured primary human pericytes to various strains of α-syn and measured if degradation was impaired in PD pericytes. Our results may inform whether targeting pericyte degradation of α-syn is a useful therapeutic option.

## MATERIAL & METHODS

### Study Design

In this study, we used different α-syn antibodies to study α-syn aggregates in pericytes derived from control and PD post-mortem brains. Pericytes were exposed for four hours to five distinct α-syn strains. α-syn degradation and abundance was measured via western blot and fluorescence microscopy for up to 21 days.

### Alpha synuclein fibril generation

Human wild-type monomeric α-syn was expressed in E. coli BL21 DE3 CodonPlus cells (Agilent Technologies) and purified as described previously [23]. To assemble human wild-type α-syn into the different fibrillar strains, the full-length monomeric protein (250µM) was incubated in 50 mM Tris–HCl, pH 7.5, 150 mM KCl for “fibrils”, in 5 mM Tris–HCl, pH 7.5 for “ribbons”, in 20mM MES pH 6.5 for “fibrils65” and in 20mM KPO4, 150mM KCl for “fibrils91” at 37°C under continuous shaking in an Eppendorf Thermomixer set at 600 r.p.m for 7 days [24,25]. A human α-syn lacking 30 C-terminal amino-acid residues, α-syn 1-110 was generated by introducing two stop codons after residue 110 by site-directed mutagenesis. This variant was purified exactly as full-length α-syn and was assembled into fibrillar structures ‘‘fibrils110’’ in 50 mM Tris–HCl, pH 7.5, 150 mM KCl. The assembly reaction was followed by withdrawing aliquots (10 µl) from the assembly reaction at different time intervals, mixing them with Thioflavin T (400µl, 10 µM final) and recording the fluorescence increase on a Cary Eclipse Fluorescence Spectrophotometer (Varian Medical Systems Inc.) using an excitation wavelength = 440 nm, an emission wavelength = 480 nm and excitation and emission slits set at 5 and 10 nm, respectively. Following assembly reaction, fibrils were fragmented to an average length of 42-52 nm by sonication for 20 min in 2 mL Eppendorf tubes in a Vial Tweeter powered by an ultrasonic processor UIS250v (250 W, 2.4 kHz; Hielscher Ultrasonic, Figure 1A). The molecular mass of fragmented fibrils was then determined by analytical ultracentrifugation [26]. Fibrils were made on average of 8300 monomers, which means that a working concentration of 2 µM equivalent monomeric α-syn corresponds to a particles (fibrils) concentration of 0.24 nM (2000/8300 = 0.24). All α-syn preparations were quantified for endotoxin levels as described previously [8,27] to prove that endotoxin levels were below 0.02 endotoxin units/mg (EU/mg) using the Pierce LAL Chromogenic Endotoxin Quantification Kit.

**Figure 1.**
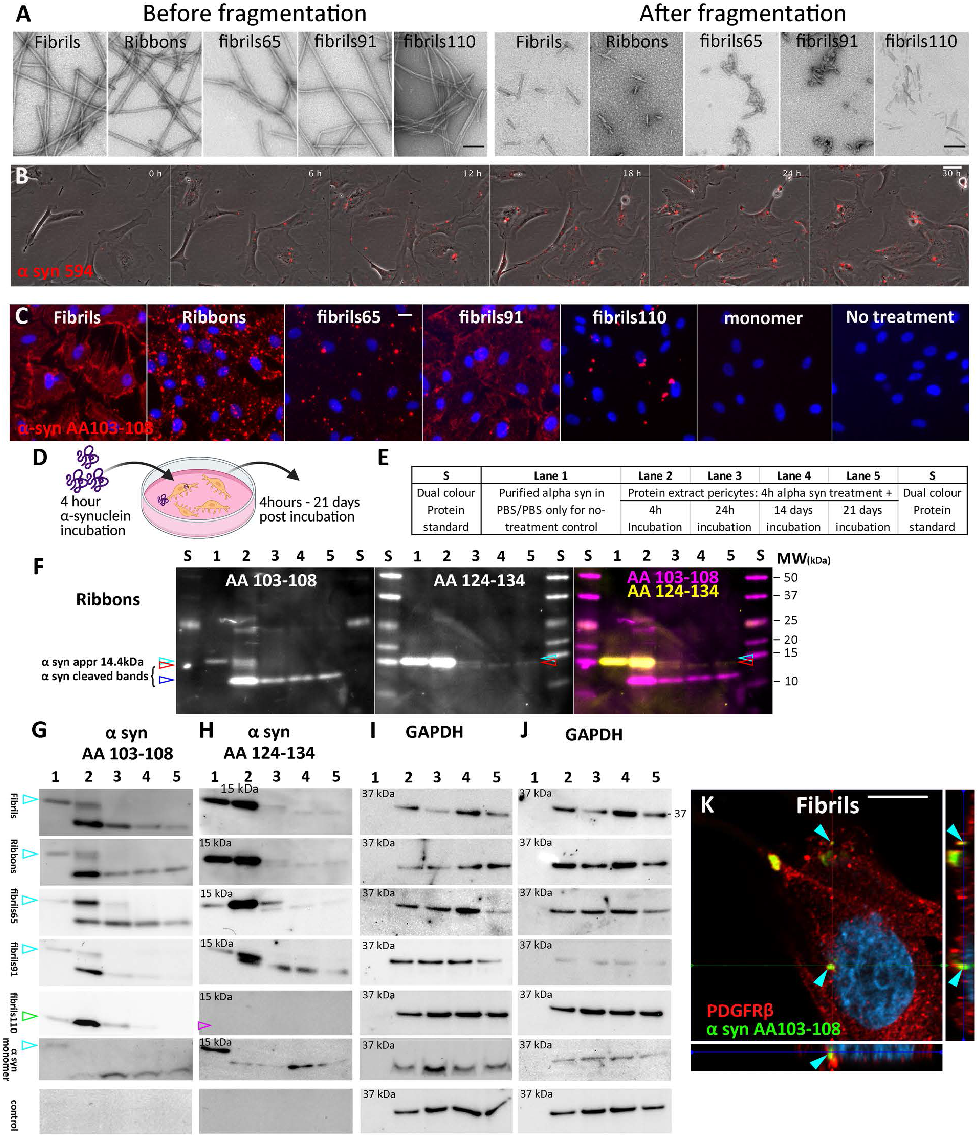
Internalisation and degradation of α-syn strains in pericytes. Electron microscopy of α-syn strains before and after fragmentation (A). Live cell imaging of fluorescently tagged α-syn for 30 hours (B). Immunofluorescent labelling of α-syn strains after 30 min in control pericytes with α-syn epitope specific antibody AA103-108 (C). Schematic representation of experimental setup (D). Loading of western blots. Lane 1 only contains pure α-syn aggregate whereas lane 2-5 shows α-syn isolated from pericytes (E). Detailed fluorescent western blot with α-syn epitope specific antibodies detecting Ribbons with α-syn antibody AA103-108 (magenta), AA124-134 (yellow) and merge showing overlap of α-syn bands. Various bands are identified indicating full length (14.4kDa, cyan arrow) and cleaved α-syn (red and blue arrows; F). α-syn detection on individual western blots for Fibrils, Ribbons, Fibrils65, Fibrils91, Fibrils110 and no-treatment control (PBS) with α-syn epitope specific antibodies showing full length α-syn (cyan arrow) and cleaved α-syn fragments. Fibrils110 lacks a full-length band as aggregate is made up of C-term cleaved α-syn (green arrow). AA 103-108 (G), AA124-134 (H), and GAPDH blot corresponds to blot shown in G after antibody stripping and relabelling (H). Full blots shown in Figure S1-S2, Confocal image with orthogonal views showing pericyte with internalised α-syn Fibrils (cyan arrows, K).

To label α-syn fibrils with extrinsic fluorophores, the fibrils were centrifuged twice at 15,000g for 10 min and re-suspended twice in PBS at 100µM, and two molar equivalents of ATTO-488 NHS-ester or ATTO-647 NHS-ester (#AD 488-35 and #AD 647-35, Atto-Tec GmbH) fluorophore in DMSO were added. The mix was incubated for 1 h at room temperature. The labelling reactions were arrested by the addition of 1mM Tris pH 7.5. The unreacted fluorophore was removed by a final cycle of two centrifugations at 15,000 g for 10 min and resuspensions of the pellets in PBS. The fibrillar nature of a-syn was assessed by Transmission Electron Microscopy (TEM) after adsorption of the fibrils onto carbon-coated 200 mesh grids and negative staining with 1% uranyl acetate using a Jeol 1400 transmission electron microscope (Figure 1).

The images were recorded with a Gatan Orius CCD camera (Gatan, Pleasanton). The resulting α-syn fibrils were fragmented by sonication for 20 min in 2 mL Eppendorf tubes in a Vial Tweeter powered by an ultrasonic processor UIS250v (250 W, 2.4 kHz; Hielscher Ultrasonic) to generate fibrillar particles with an average size 42-52 nm as assessed by TEM analysis.

### Human brain tissue

The post-mortem adult human brain tissue for this study was obtained from the Neurological Foundation Human Brain Bank (Centre for Brain Research, University of Auckland). The University of Auckland Human Participants Ethics Committee approved the protocols used in these studies (approval# 011654). The Health and Disabilities Ethics Committee New Zealand approved the use of surgical removed brain tissue (biopsy tissue approval# AKL/88/025). Written, informed consent was obtained from the individual(s) and/or next of kin to use the tissue. Pathological assessment of neurologically normal cases (controls) reported no disease-related pathology beyond that expected in aged individuals, and all PD cases were confirmed by a neuropathologist (Table 1). The neurosurgical brain tissue was obtained with written informed consent from Auckland City Hospital under HDEC ethics approval AKL/88/025.

**Table 1.**
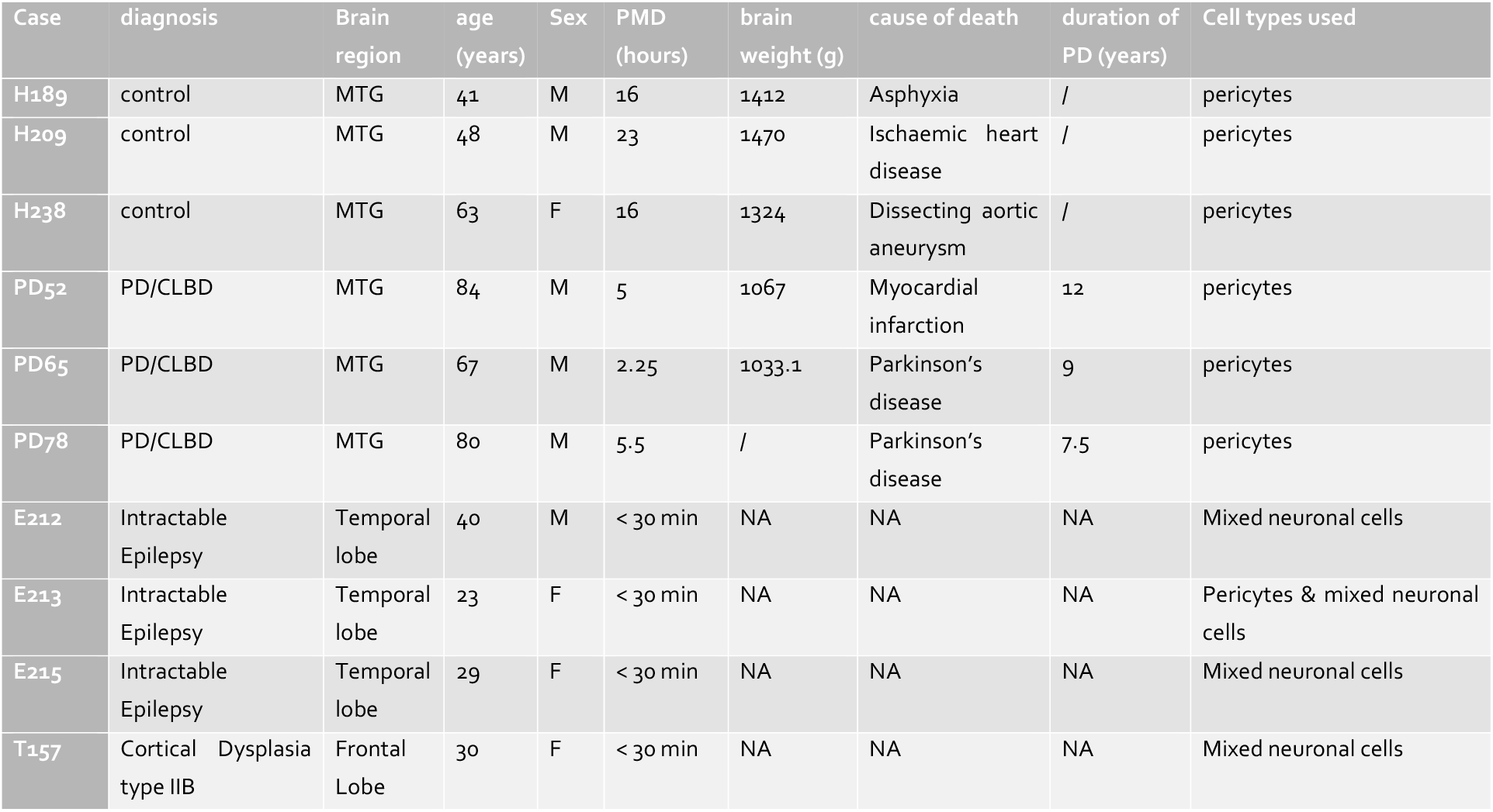
Case information for post-mortem and surgical removedbrain tissue used in this study.

| Case | diagnosis | Brain region | age (years) | Sex | PMD (hours) | brain weight (g) | cause of death | duration of PD (years) | Cell types used |
| --- | --- | --- | --- | --- | --- | --- | --- | --- | --- |
| H189 | control | MTG | 41 | M | 16 | 1412 | Asphyxia | / | pericytes |
| H209 | control | MTG | 48 | M | 23 | 1470 | Ischaemic heart disease | / | pericytes |
| H238 | control | MTG | 63 | F | 16 | 1324 | Dissecting aortic aneurysm | / | pericytes |
| PD52 | PD/CLBD | MTG | 84 | M | 5 | 1067 | Myocardial infarction | 12 | pericytes |
| PD65 | PD/CLBD | MTG | 67 | M | 2.25 | 1033.1 | Parkinson's disease | 9 | pericytes |
| PD78 | PD/CLBD | MTG | 80 | M | 5.5 | / | Parkinson's disease | 7.5 | pericytes |
| E212 | Intractable Epilepsy | Temporal lobe | 40 | M | < 30 min | NA | NA | NA | Mixed neuronal cells |
| E213 | Intractable Epilepsy | Temporal lobe | 23 | F | < 30 min | NA | NA | NA | Pericytes & mixed neuronal cells |
| E215 | Intractable Epilepsy | Temporal lobe | 29 | F | < 30 min | NA | NA | NA | Mixed neuronal cells |
| T157 | Cortical Dysplasia type IIB | Frontal Lobe | 30 | F | < 30 min | NA | NA | NA | Mixed neuronal cells |

### Isolation of primary human brain pericytes

Middle temporal gyrus (MTG) tissue was used to generate pericyte cultures from human post-mortem brains with a PD diagnosis or pathologically normal (control) cases as determined by a neuropathologist (Table 1). We found that post-mortem delay (PMD) had less effect on the viability of pericyte cultures compared to the age at death and disease diagnosis. Viable pericytes can be grown from brain tissue up to 48 hours post-mortem. A detailed description for generating pericyte cultures by our group has previously been described [28]. In short MTG tissue was mechanically dissected and dissociated before enzymatic digestion in Hank’s balanced salt solution (HBSS), containing 2.5 U/mL papain (Worthington) and 100 U/mL DNase 1 (ThermoFisher) for 30 minutes at 37°C with gentle rotation. This step included a gentle titration at 15 minutes. Following this, complete media, DMEM: F12 (ThermoFisher), was used to stop the enzymatic digestion. To collect the cells, they were centrifuged (170g, 10 min) and re-suspended in complete media. Cells were plated onto uncoated T75 culture flasks (Nunc). Cells were subsequently incubated at 37°C with 5% carbon dioxide until seeded for experiments, grown in DMEM:F12 (v) containing 10% fetal bovine serum (ThermoFisher) and 1% Penicillin/Streptomycin (ThermoFisher). For all the experiments presented here, pericytes were at passages 5-9. This ensures pericyte pure cultures without contamination of astrocytes or microglia. Late passages (> P4) displayed immunocytochemical staining for pericyte markers including PDGFRβ, alpha-smooth muscle actin (αSMA), neural/glial antigen 2 (NG2), CD146 and desmin. The cultures also displayed immunocytochemical staining for fibroblast markers prolyl-4-hydroxylase (P4H) and fibronectin [1,4]. Pericytes grown in these culture conditions have been well characterized, carry a unique inflammatory profile, and can phagocytose large beads [1,2,4,7,9,12,14].

### Cell plating and incubation with α-syn aggregates

Cells were harvested for experiments by adding 2 mL 0.25% Trypsin-1mM ethylenediaminetetraacetic acid (EDTA; ThermoFisher) and incubated for 2-5 minutes at 37°C to allow for cell detachment. Cells were collected in warm DMEM: F12 media. 10 μL of 1:1 Trypan Blue (ThermoFisher) and the cell suspension was prepared and added to a haemocytometer for cell counting. Cells were re-suspended in the correct volume of DMEM: F12 to achieve the required cell density of 5,000 cells/well in 96-well plates (Nunc) or 50,000 cells in 6 well plates. Pericytes were treated at 37°C with the various α-syn strains (Fibrils, Ribbons, fibrils65, fibrils91 or C-terminus truncated fibrils110, 100 nM in PBS) or PBS (no treatment control) for 4 hours, washed and subsequently returned at 37°C until endpoint (0 hours – 37 days). Results up to 21 days are shown in this manuscript as no distinctive changes were observed between 21-37 days.

### Primary mixed neuronal cultures

Grey matter isolated from normal cortex was mechanically dissociated into <1 mm pieces, followed by enzymatic digestion. Cells were grown in neuronal growth medium (DMEM: F12, supplemented with 2% B27, Penicillin/Streptomycin, GlutaMAX®, (ThermoFisher) 10 µM Y-27632 2HCl (Selleckchem); 2 µg Heparin (Sigma), 40 ng/ml of NGF, BDNF, NT-3, GDNF, and IGF-1 (Peprotech), plated onto poly-D-lysine (100µg/ml) coated coverslips and incubated at 37 °C with 5% CO_2_. The culture media was half changed every 24 hours for the first 2 days and every 3-4 days after that. These cultures are extensively characterised and contain a mixture of neurons with mature neurophysiological properties, astrocytes, pericytes and microglia [33].

### Live-cell Imaging

Pericytes, cultured in small Petri dishes (150318, Nunc, ThermoFisher), were treated with 100nM α-syn-atto 647 for 4h, washed and placed in a Nikon Biostation cell incubator (5% CO_2_, 37°C, Plan Fluor 20x/ 0.5 NA DL) for up to 120 h. Images of brain pericytes were captured every 5 min during this period. During imaging, no readily identifiable adverse phototoxic effects were observed on the cells.

### Protein isolation and western blotting

Whole pericytes cell lysate was rinsed with ice-cold PBS and isolated at the endpoint from 6 well plates with 100µl sample harvesting buffer (62.5 mM Tris-HCl pH 6.8, 2% SDS, 10% glycerol). Cells were scraped off, boiled at 100°C for 10 min, and the lysate was stored at-80°C. Western blot was performed according to the manufacturer guidelines using Novex 10-20% Tris-Glycine Mini Gels (XP10202BOX, ThermoFisher). Proteins were subsequently electrotransferred onto a polyvinylidene difluoride membrane (PVDF, Hybond-P; Amersham) using Novex Tris-Glycine Native Running Buffer in a BioRad blot module. For immuno-staining, the membrane was blocked for one hour using Odyssey blocking buffer/TBS-T (1:1; 0.1% Triton-X). Primary antibodies were incubated overnight at 4°C in Odyssey blocking buffer/TBS-T (1:1). Antibodies raised against α-syn C-terminus (Mouse anti α-syn, Amino Acid (AA) 124-134 (1:1,000) and ab1903 (Abcam); Mouse anti α-syn 4B12, AA 103-108, IgG1 (1:2,000, MA1-90346, ThermoFisher), GAPDH (1:1,000, ab9484, Abcam). Blots were washed in TBS-T and incubated with fluorescently conjugated secondary antibodies for 3 hours at room temperature in Odyssey blocking buffer/TBS-T (1:1 and 0.02% SDS; IRDye 800CW Donkey anti-mouse, 926-32212 (Li-COR)). Blots were washed (3 × 5min TBS-T and 2 × 5 min TBS) before being imaged on the ChemiDoc MP Imaging System (Biorad). Subsequently, gels were stripped for 30 min at 60°C (62.5 nM Tris, pH6.8; 2%SDS; 100 mM β-mercaptoethanol), extensively washed in TBS-T before being re-probed for GAPDH. For these experiments, the samples were run on two gels, with the first blot labelled with AA103-108 and the second blot labelled with AA124-134. After imaging both gels, they were stripped and relabelled for GAPDH. Full western blots are available in supplementary information (Figure S 1 Figure S 2).

### Immunocytochemistry

At endpoints, cells were fixed in 4% paraformaldehyde for 15 min and washed/permeabilised with phosphate buffered saline (PBS) with 0.2% Triton X-100TM (PBS-T). Cells were incubated overnight at 4 °C with primary antibodies — targeting α-syn (Mouse anti α-syn, AA 124-134, ab1903, Abcam; Mouse anti α-syn 4B12, AA 103-108, IgG1, MA1-90346, ThermoFisher; pericytes (Rabbit anti-PDGFR β, ab32570, Abcam; gt PDGFR β, AF-385, R&D systems) — diluted in immunobuffer (1% goat or donkey serum, 0.2% Triton X-100 in PBS). Cells were rewashed in PBS-T and incubated with filtered (0.22 µm syringe filter, Merck) fluorescently conjugated secondary antibodies and 20 μM Hoechst 33258 (ThermoFisher) for 2 hours at room temperature. Images were acquired using the automated fluorescence microscope ImageXpress® Micro XLS (Version 5.3.0.1, Molecular Devices) using the 20 x (0.45 NA) CFI Super Plan Fluor ELWD ADM objective lens and Lumencor Spectra X configurable light engine source. Confocal images were acquired using a LSM 800 Airyscan confocal microscope (Zeiss LSM 800 Airyscan confocal microscope) with a 40x/1.3 or 63x/1.4 NA Plan Apochromat oil immersion using the Airyscan module.

### High content image analysis

Images acquired on the ImageXpress® Micro XLS at respective endpoints were exported from MetaXpress (software information) as uncompressed 16-bit TIFFs for analysis using FIJI (version #1.52p). A rolling-ball median filter (radius = 2.0 pixels) was applied to Hoechst images for background subtraction before running “Find Maxima” to define cell boundaries using Voronoi tessellation (prominence = 500). Cell boundaries were then added to the region of interest (ROI) manager, excluding cells on edges of image frames. Cell nuclei were then auto-threshold masked using the ‘Huang’ setting to find the cell nucleus area. Next, the α-syn images (AA103-108) were processed by applying a median (radius = 2.0) and Gaussian filter (sigma = 15.0) to duplicates of the image before subtracting the gaussian processed image from the median processed image to reduce noise and identify background, respectively. The resulting images were thresholded (82, 65535) using the ‘Default’ method before converting to a binary mask. These masks were then counted, with a minimum aggregate size of 2 pixels^2^. For intensity measurements, intensity measurements within cell masks were made on raw aggregate images. This process was batch processed on the entire image set using a custom ImageJ macro. The script for this analysis is available in supplementary information.

### Immunohistochemistry

For the immunohistochemical studies, the human brains were processed and labelled as previously described [22,34,35]. 7 µm-thick sections from paraffin-embedded MTG blocks were cut using a rotary microtome (Leica Biosystems RM2235) and mounted onto Über plus printer slides (InstrumeC) in a water bath set to 42°C (Leica Biosystems H1210). Sections were left to dry for 72 hours at room temperature prior to immunohistochemical staining.

Sections were then blocked in 10% normal goat serum in PBS for 1 hour at RT. Subsequently, the sections were incubated with primary antibodies (α-synuclein-phospho S129, ab51253, Abcam) overnight at 4°C. Sections were incubated with the corresponding goat secondary Alexa Fluor (488, 594, 647) conjugated secondary antibody (ThermoFisher) and Hoechst 33342 (Thermofisher) at 1:500 for 3 hours at RT. Control sections where the primary antibody was omitted showed no immunoreactivity. The control experiments showed that the secondary antibodies displayed no cross-reactivity. Fluorescent images were captured with a MetaSystems VSlide slide scanner with a 20x dry lens (NA 0.9).

### Data visualisation and statistical hypothesis

Data visualisation and statistical hypothesis testing were performed using GraphPad Prism® Version 9.00 and R Version 4.0. The colours represent a 2D kernel density estimation. It is scaled to 1 for each graph, with the bright yellow area displaying the highest density of cells. All experiments were performed in at least three independent cases. In general, data are expressed as individual cells or average/case. The PMD and age of death in PD vs control brain donor were different (control PMD:18.4 ± 4.0, age: 50.6 ±11.2; PD PMD: 4.3 ± 1.8, age: 77 ± 8.9). For this reason, PD and control pericytes (n = 3 each group) were analysed separately. Data visualisation and statistical hypothesis testing were performed using GraphPad Prism® Version 7.00. Two-way analysis of variance (ANOVA) was used when comparing responses across a number of stimuli with Tukey’s post hoc adjustment for multiple comparisons. Linear regression was used to analyse correlations. One-way analysis of variance (ANOVA) was used when comparing across cell types with Tukey’s multiple comparison adjustment. Statistical significance was set at p < 0.05. Final figures were composed using Adobe Photoshop CC (Adobe Systems Incorporated, v20.0.6).

## RESULTS

### α-syn aggregates are internalised and cleaved by pericytes

Live cell imaging showed rapid uptake of various fibrillar α-syn strains (Fibrils, Ribbons, fibrils65, fibrils91 or fibrils110) in pericytes when added to the culture media. α-syn formed discrete spots within the pericytes (Figure 1B) in agreement with what we reported using astrocytes induced from human pluripotent stem cells [16]. Binding and internalisation of fibrillar α-syn was strain-dependent, with apparent differences observed within 30 min after treatment in agreement with what we reported for rodent neurons [36]. Fibrils and fibrils91 are primarily found membrane bound on the pericytes. As uptake progressed, this distribution gradually changed to a punctate pattern, whereas Ribbons, fibrils65 and fibrils110 already showed a punctate pattern. 30 min after adding fibrillar α-syn, all cells within the culture displayed positive α-syn labelling except fibrils110, where only a subset of cells contained α-syn. No puncta were observed in pericytes treated with monomeric α-syn (Figure 1C; Figure S 3B).

Pericytes were subsequently incubated with each of the five α-syn strains (Fibrils, Ribbons, fibrils65, fibrils91 or fibrils110) to test their ability to degrade α-syn (experimental setup, Figure 1D). Pericytes were pre-treated with fibrillar α-syn for 4 hours before excess α-syn was washed off and followed for an additional 4 hours - 21 days Two antibodies directed against the C-terminal part of α-syn (AA103-108 and AA124-134) clearly labelled exogenous α-syn (before addition to cells). For western blot gel layouts refer to Figure 1E. Protein extracts from pericytes exposed to α-syn (Ribbons) showed immunoreactivity at approximately 14 kDa and two lower bands bearing AA103-108 (white, Figure 1F); and 14kDa and one lower band bearing AA124-134 (magenta; Figure 1F). Full length α-syn (14 kDa) was no longer visible after 14 days. On the contrary cleaved α-syn bands remained visible up to 21 days with AA103-108 and AA124-134.

When comparing the five α-syn strains, we observed a similar pattern for Fibrils, Ribbons, fibrils65 and fibrils91 (Figure 1G-H; GAPDH bands for each blot Figure 1I-J). Full length α-syn was observed up to 24 hours (cyan arrow) and cleaved α-syn was seen until 21 days. Because fibrils110 is comprised of C-terminus truncated protein, we only observed a lower α-syn band (green arrow) corresponding to the cleaved fragment observed in the other strains when labelled with AA103-108 (Figure 1G). AA103-108 detected fibrils110 (Figure 1G), whereas AA124-134 did not (magenta arrow, Figure 1IH), thereby confirming that the C-terminus portion was lacking from fibrils110. No α-syn bands were present in the untreated control (bottom panel: Figure 1G-I). These results indicate that pericytes have low or do not have endogenous α-syn, but they can internalise and process α-syn aggregates into discretely sized fragments. Full length α-syn could not be detected by western blotting after 24 hours, whereas cleaved α-syn remained present within pericytes for up to 21 days. These findings show that AA103-108 and AA124-134 detect full length α-syn. Two different α-syn bands are identified by AA103-108 and AA124-134, indicating that pericytes cleave α-syn at two different cleavage sites. AA103-108 was the only antibody that consistently identified cleaved α-syn for all strains.

α-syn was immunolabelled within pericytes using the AA103-108 antibody that enables detection of internalised full length and cleaved fibrillar α-syn (cyan arrows show internalised α-syn fibrils; Figure 1K). Monomeric α-syn was not included in this time course as no α-syn puncta were observed (Figure S 3B-C).

When qualitatively comparing uptake five hours after treatment, Fibrils, Ribbons, and fibrils91 showed a similar pattern of punctate aggregates of different sizes (Figure 2 A-F). Fibrils65 had a similar pattern, with fewer overall aggregates identified. However, larger aggregates were occasionally visualised (magenta arrow). The biggest differences were observed between fibrils110 and the other strains. Fibrils110 had a low number of highly immunoreactive aggregates per cell, with several cells devoid of aggregates (white arrow). Despite the large variability observed between cells treated with different strains, some general observations were apparent. Fibrils, Ribbons and fibrils91 displayed immunoreactivity for many small aggregates in most cells whereas pericytes treated with fibrils110 displayed a small number of large immunoreactive aggregates in a small proportion of cells (Figure 2 A-F). This agrees with previous observations we made using these strains and astrocytes induced from human pluripotent stem cells [16]. No phosphorylated α-syn was observed at any of the timepoints (Figure S 3B-C).

**Figure 2.**
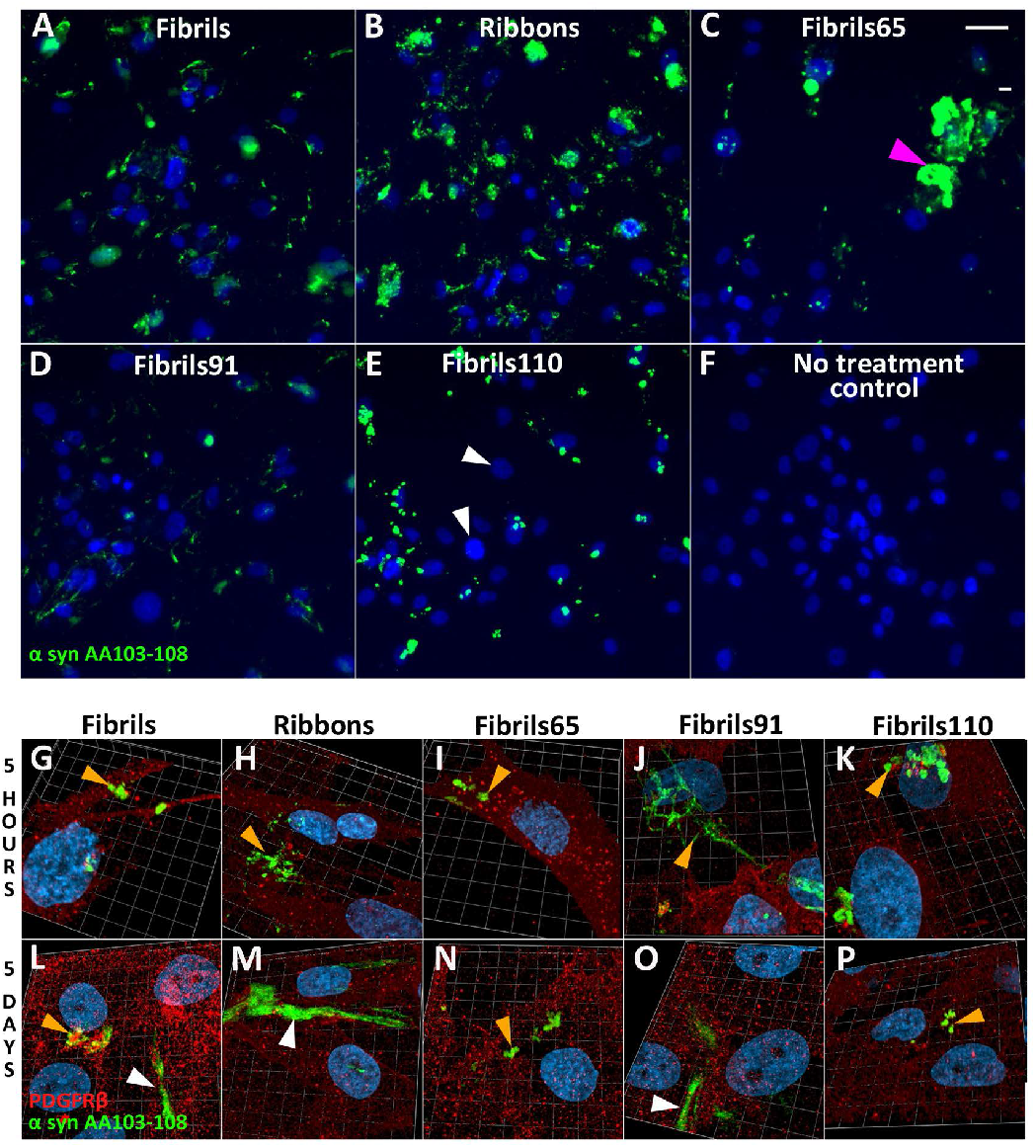
Immunofluorescent labelling of α-syn strains in control pericytes with α-syn epitope specific antibodies AA103-108 (green). Strains specific fluorescent staining 5 hours after pre-treatment with 100nM α-syn. At early timepoint all cells contain aggregates, except for fibrils110 treated cells (white arrows); Fibrils (A), Ribbons (B), fibrils65, with occasional larger aggregates (magenta arrow) (C), fibrils91(D), fibrils110 (E), no treatment control (F). 3D confocal view of α-syn aggregates in pericytes 5 hours and 5 days after pre-treatment with 100nM Fibrils (D, I), Ribbons (E, J), fibrils65 (F, J), fibrils91 (G,K), fibrils110 (H,L). Typically aggregates appear as spots (orange arrows) with thicker fibres (white arrows) appearing more frequently at later timepoints. Scale bars represent 50 µm.

Live cell imaging of all strains further confirmed these observations. Throughout the recordings, pericytes frequently underwent mitosis, indicating that internalised fluorescently tagged α-syn aggregates do not inhibit cell division (Movie S1-5).

3D Confocal Airyscan images of pericytes labelled with PDGFRβ (red) treated with α-syn aggregates confirmed previous observations at 5 hours with AA103-108. After 5 days α-syn was present in the form of punctate aggregates (orange arrows) and elongated fibres (white arrows; Figure 2G-P).

### Clearance of aggregated α-syn in pericytes derived from control and PD brains is dependent on the strain and exposure time

Following pre-treatment with α-syn aggregates (4 hours, 100nM), pericytes fixed at 7 time-points (0-21 days), were labelled with AA103-108 antibody. α-syn aggregate density gradually decreased over time with α-syn AA103-108 remaining visible up to 21 days. α-syn was present, in the form of punctate aggregates and elongated fibres (green arrows, Figure 3). Fibrils91 staining decreased faster than Fibrils, Ribbons and fibrils65, with clear differences observed after 1 day. As described above, fibrils65 occasionally displayed large, intense α-syn aggregates (magenta arrow). These large fibrils65 puncta are a visualisation artifact caused by representing all the strains with the same brightness settings. Subsequent analysis performed on the raw data showed that these large fibrils65 puncta are comprised of several individual aggregates. The untreated control showed no α-syn immunolabelling (Figure 3). This labelling validated our western blot data while including additional timepoints.

**Figure 3.**
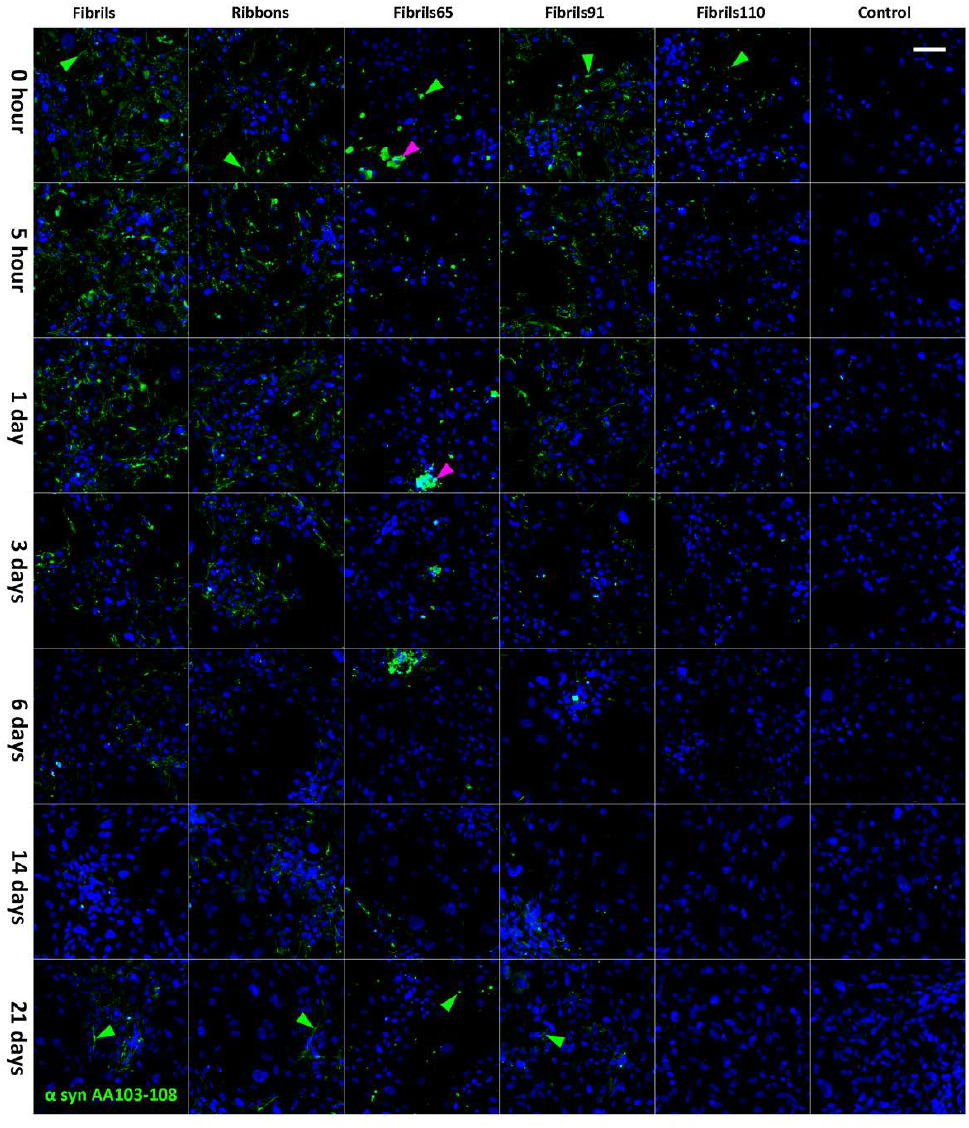
Immunofluorescent labelling of α-syn strains in pericytes with α-syn specific antibodies AA103-108 (green) after pre-treatment with 100nM Fibrils, Ribbons, fibrils65, fibrils91 and fibrils110. Large aggregates were occasionally found for fibrils65 (magenta arrow). Scale bar represent 100 µm

To assess α-syn strain clearance in pericytes, automated quantification of aggregated α-syn detected with AA103-108 antibodies in single cells was conducted (Figure 4). Control (n=3) and PD derived pericytes (n=3) were analysed separately for this study. Individual pericytes revealed variation in the average aggregate size/cell and average aggregate count/cell for each strain. Nevertheless, mean values for each case are representative of our observations.

**Figure 4.**
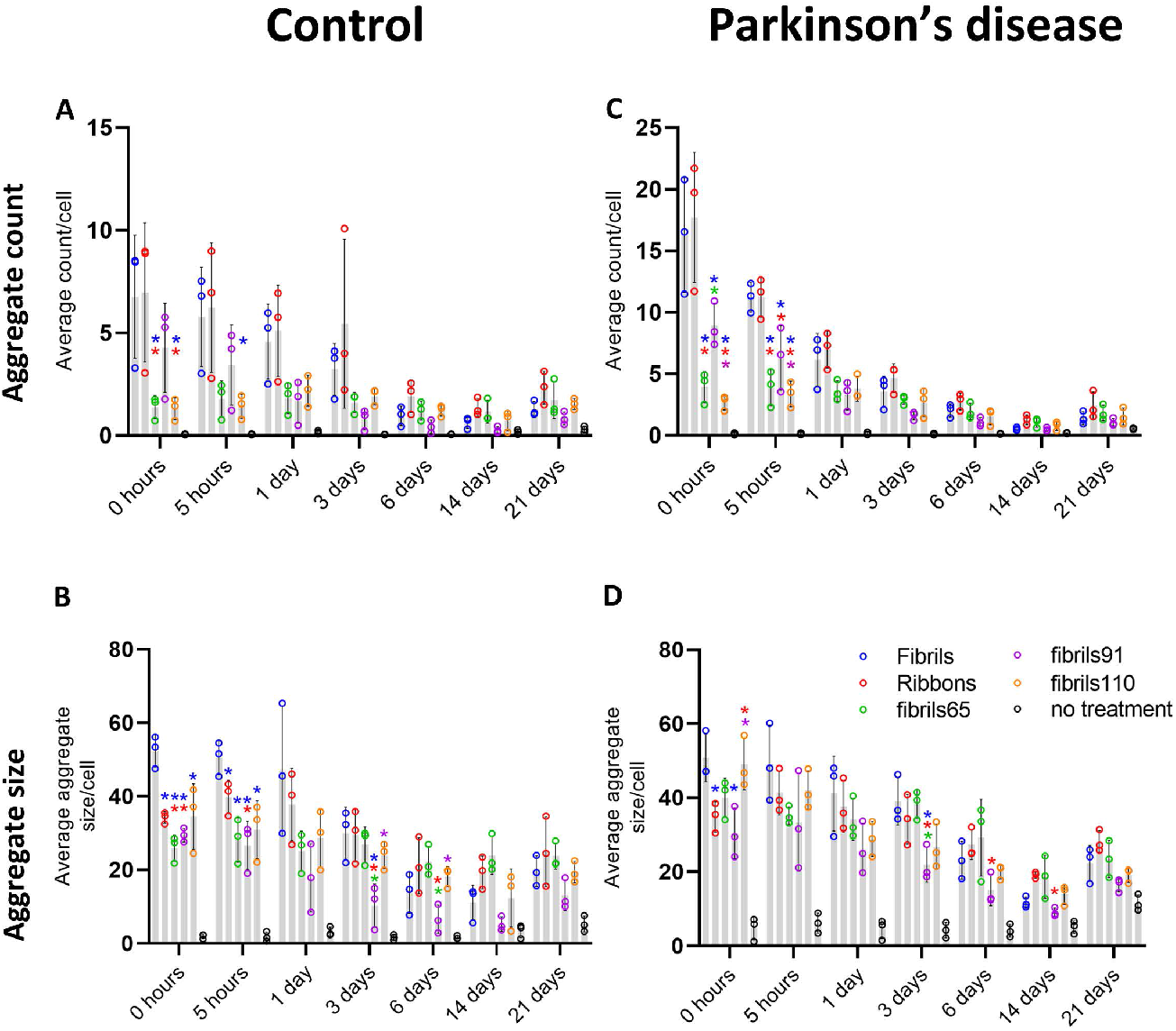
Strain specific differences in control and PD derived post-mortem pericytes for all strains. Aggregate count/cell analysis and average aggregate size/cell analysis based on AA103-108 for control pericytes (A-B), and PD pericytes (C-D) significance indicated by coloured * (p<0.05). blue * indicates significant difference with Fibrils; red * indicates significant difference with Ribbons, green * indicates significant difference with fibrils65; purple * indicates significant difference with fibrils91.

At 0 hours, there were significantly more aggregates/cell for Fibrils and Ribbons compared to fibrils65 and Fibrils110 in both control and PD pericytes. This difference remained up to the 5 hour time point in the PD pericytes. There were also more fibrils91 aggregates in PD pericytes than fibrils65 and Fibrils110. This is most likely due to the increased amount of aggregates/cell observed for Fibrils, Ribbons and fibrils 91 in PD pericytes. In contrast, the amount of aggregates/cell remained similar for fibrils65 and fibrils110 in control and PD pericytes. The average aggregate size for Fibrils was higher at 0 and 5 hours compared to all other strains, but this was not the case in PD pericytes, where Fibrils were only larger than Ribbons and P91. Over the first three days there was a decline in the size and number of aggregates for all strains except for fibrils 65 and fibrils110. Fibrils65 and fibrils110 displayed lower aggregate counts from the beginning, and these counts remained more constant until six days. Fibrils91 aggregates appeared to reduce faster in size compared to the other strains and reached significance after three days (Figure 4 A-D). Overall, both average counts/cell and aggregate size/cell decreased over time, indicating that the fewer remaining aggregates were also smaller. Aggregates remained visible at the latest timepoints with many pericytes still containing aggregated α-syn 21 days after exposure. (Figure 4 A-D).

Combined representation of aggregate counts and average aggregate size for each cell in the form of density plots showed a greater variability in PD-derived compared to control pericytes for all strains. The vertical bars represent the fraction of cells with aggregates (Figure 5). For Ribbons, most of the control pericytes contained a low number of small aggregates (up to 10 counts/cell, colours yellow to purple on the plot). The aggregate counts spread in PD-derived pericytes went up to 28 counts/cell while maintaining the same aggregate size range. Immediately after exposing pericytes to Ribbons, at time 0h, the proportion of pericytes containing aggregates from control and PD was 94.4 and 99.5%, respectively. With time, at day 14, the proportion of pericytes containing aggregates from control and PD decreased to 57.5 and 59.3%, respectively indicating that a high proportion of pericytes still contained α-syn aggregates (Figure 5A). For fibrils110 control pericytes displayed limited variation with most cells containing 1-5 aggregates while that varied more in PD pericytes (up to 10 counts/cell). The number of cells containing aggregates differed markedly upon exposing pericytes to fibrils110. A large proportion of control pericytes were devoid of aggregates at 0h (43% for controls, 11.7% for PD-derived pericytes). With time, at day 14, the proportion of control and PD-derived cells without aggregates increased (53.3% and 53.6%, respectively; Figure 5 B).

**Figure 5.**
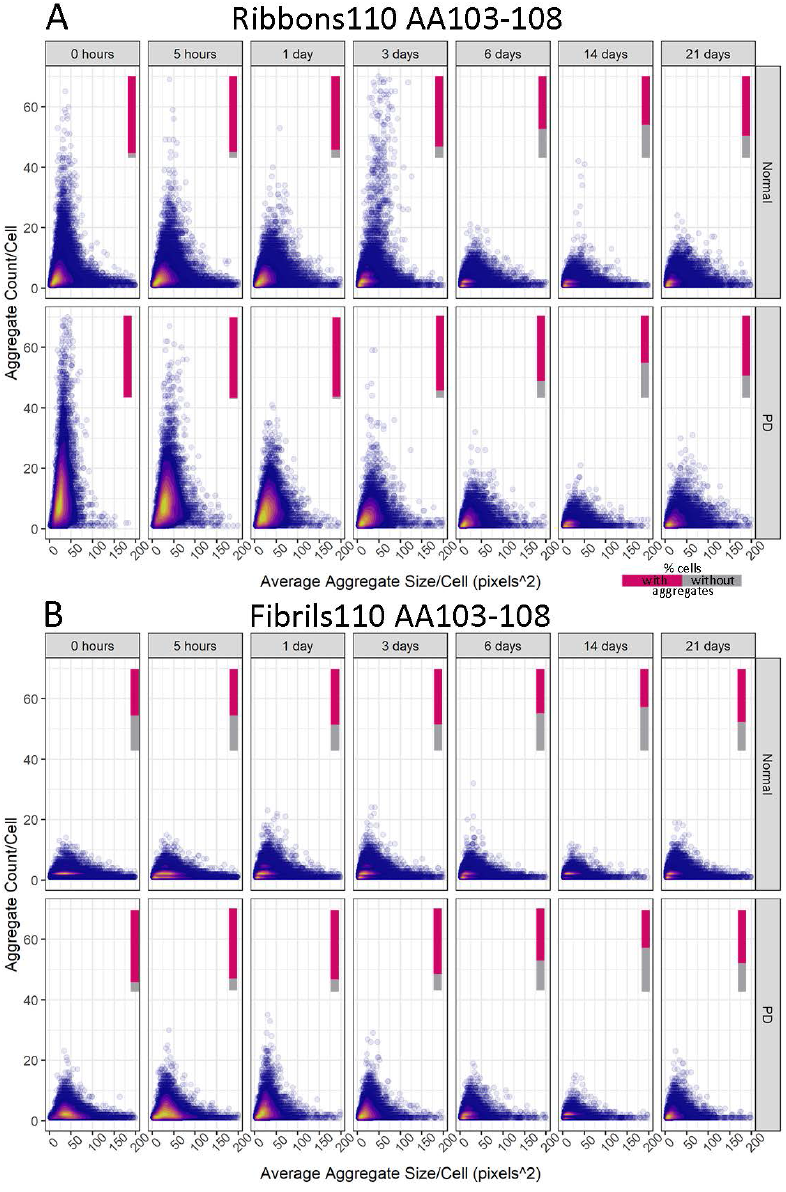
Density plots showing a differential response to α-syn aggregates for control and PD-derived pericytes based on AA103-108 for Ribbons and fibrils 110. Cells without aggregates are excluded from density plots. Relative amounts of cells with aggregates represented in bar (% cells with aggregates in magenta, % cells without aggregates in grey).

The density plots observed for Fibrils and fibrils91 followed that of Ribbons, whereas fibrils65 was similar to fibrils110. Fluorescent debris was picked up in the untreated control and gradually increased with time (0h-14 days; 5 to 10% of control, 15-16% PD-derived pericytes) and peaked at 21 days (20% and 36%, respectively) (Figure S 4A-D).

### Sustained α-syn internalisation by pericytes in human primary mixed neuronal cultures

Differences may exist between uptake and clearance of α-syn in a monoculture and mixed cell populations. Indeed, pericytes might no longer internalise α-syn aggregate in the presence of other cell types. To show uptake, we added fluorescently labelled α-syn Fibrils for 60 hours to primary cell cultures consisting of mature human primary neurons, astrocytes, microglia (red arrow) and pericytes (yellow arrow). The neurochemical phenotype and functionality of these neurons have been extensively validated [28,33]. The average age of donors for the mixed neuronal cell cultures in this study was (30.5 ± 7 years). One cell type displayed intense labelling (red) indicative of rapid binding and internalisation of α-syn Fibrils from the culture media (red arrow, Figure 6A, Movie S 6). These cells are CD45^+^/Iba1^+^ microglia that rapidly internalised α-syn (red arrows Figure 6 B-G). The proximity of microglia did not prevent α-syn fibrils internalisation by PDGFRβ^+^ pericytes (yellow arrow, Figure 6H, Movie S 6). Within the timeframe (96 hours), MAP2^+^ neurons (blue arrow, Figure 6I), internalised α-syn Fibrils to a much lesser extent. Confocal imaging with orthogonal projection demonstrates the difference in α-syn fibrils internalisation between microglia (red arrow) and neurons (blue arrow) visible (Figure 6J). These co-culture experiments revealed that pericytes in mixed cell populations retain their capacity to internalise fibrillar α-syn aggregates.

**Figure 6.**
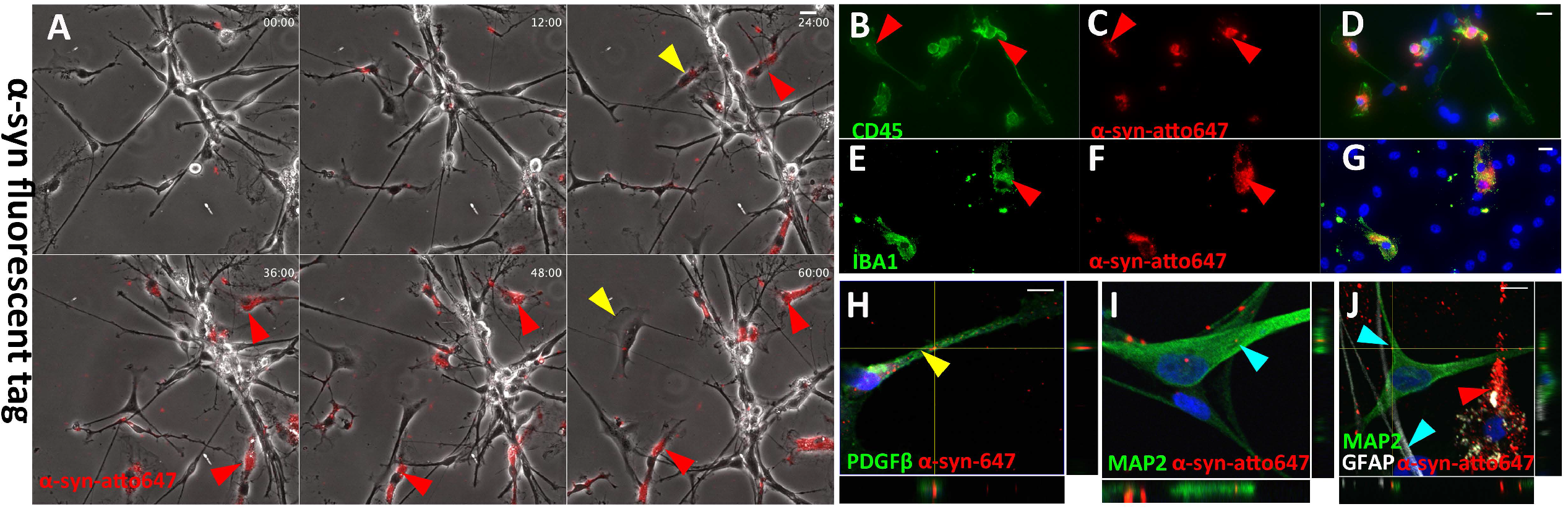
Primary human mixed neuronal cell exposed to α-syn aggregates - internalisation is cell type specific. Representative images of live cell recording with α-syn-594 aggregates(A). Confocal imaging indicating that α-syn-594 aggregates are present within neurons and pericytes (B-C). Cells with high α-syn aggregate load are microglia CD45^+^ and IBA1^+^ (E-I). Representative images of primary human mixed neuronal cell exposed to Ribbons (J) with Ribbons within Astrocytes GFAP high, MAP2 low (L-O) and neurons GFAP low-MAP2 high (P-S)

## DISCUSSION

Situated at the blood and the brain parenchyma interface, pericytes play an essential role in regulating blood-brain barrier function and as mediators of neuroinflammation [17,37]. In PD, decreased pericyte coverage of vessels is coupled with increased leakiness of the blood-brain barrier [20,21]. Our previous work showed that in situ human brain pericytes contain α-syn aggregates in similar proportions to microglia and astrocytes [13]. Furthermore, primary human pericytes can transfer α-syn through nanotubes in vitro to neighbouring cells [22]. It remains unclear whether pericytes function as a mere conduit for α-syn spread or if they are also involved in aggregate seeding. Pericytes have either low or no endogenous expression of α-syn, suggesting seeding in a pure pericyte culture is unlikely. We, therefore, hypothesise that pericytes must internalise α-syn if they are involved in aggregate seeding. Here, we demonstrate that pericytes efficiently internalise fibrillar α-syn and efficiently break down α-syn aggregates in vitro. Uptake and clearance kinetics differ when comparing five pure α-syn strains (Fibrils, Ribbons, fibrils65, fibrils91 and fibrils110), with Ribbons being the most efficiently taken up and fibrils91 cleared the fastest. As such, pericytes could − together with microglia and astrocytes − act as barriers that remove α-syn aggregates, thereby reducing α-syn induced toxicity and slowing down spread, in turn delaying disease onset.

It is well-established that in synucleinopathies Lewy body pathology spreads throughout the central nervous system in disease specific patterns [4]. α-syn is a constitutive component of Lewy bodies and neurites and is known to aggregate into large fibrillar assemblies [38]. Once a cell contains aberrant α-syn, there are three potential outcomes: degradation, deposition in a specialised compartment (inclusion), or release into the extracellular space. Upon release, α-syn can be taken up by acceptor cells where the cycle of protein aggregation in a prion-like manner continues, or degradation occurs. Unlike pericytes, astrocytes and microglia are well studied in this aspect. Different brain cells take up exogenous α-syn with different kinetics [39]. Microglia are more efficient than neurons and astrocytes in both uptake and degradation of extracellular α-syn [39,40]. This suggests that the uptake pathways are differentially regulated or that different cell types are equipped with distinct receptors for extracellular α-syn [40]. As such interaction with fibrillar α-syn is likely influenced in mixed cell cultures. Therefore, we demonstrate that pericytes internalise α-syn both in monoculture and when combined with other cell types known to phagocytose α-syn [14,39,41]. In both culture conditions, pericytes efficiently take up fibrillar α-syn. Uptake by pericytes still occurred despite the proximity of professional phagocytes like microglia, strengthening the results obtained from our pericyte monoculture.

Specific cleavage of α-syn by pericytes has not been previously studied. In this study, using different epitope-specific α-syn antibodies directed at the α-syn C-terminus we show that human primary pericytes can efficiently degrade α-syn aggregates through cleavage into several smaller fragments. These fragments correspond to what has previously been observed in the BV-2 microglial cell line, primary murine microglia, astrocytes and SH-SY5Y cells [14,39,42]. Truncated α-syn is consistently found in human brain tissue; AA1-119 (truncated at Asp-119) and AA1-122 (truncated at Asn-122) have an abundance as high as 20-25% relative to full-length α-syn [43,44]. We did not assess the exact degradation mechanism of fibrillar α-syn by pericytes in this study. However, pericytes express high levels of lysosomal granules [16–18]. Thus, degradation of α-syn can occur both via autophagic clearance and by proteasomal degradation. It is known that the dysfunction of both systems is linked to PD pathogenesis [19–22]. Our current research is pointing towards lysosomes being primarily responsible for α-syn degradation in pericytes.

In this study, we further determined that the cleavage and ongoing detection of fibrillar α-syn in pericytes are strain-dependent. Overall, distinct strains displayed different aggregate count kinetics, with Fibrils and Ribbons having high aggregate counts. They are closely followed by fibrils91, whereas fibrils65 and fibrils110 have a limited number of aggregates per cell. We show that all strains undergo degradation but that cleaved α-syn species remain present for at least 21 days. Our findings suggest that Fibrils and Ribbons remain in pericytes longer compared to other strains, potentially indicating that pericytes have difficulty removing these strains. Based on our data, we speculate that fibrils65 and fibrils110 are internalized at a slower rate, which would explain the lower number of aggregates/cell for these strains. On the other hand, fibrils91, which has higher aggregates/cell than fibrils65 and fibrils110, is reduced in size more than the other strains, which points to faster degradation. This indicates a difference in processing. It is hard to speculate the exact contribution of internalization versus processing for each strain. Still, both likely play a role in the clearance of fibrillar α-syn by pericytes.

A direct comparison between pericytes derived from control and PD post-mortem brain was not possible due to differences in PMD and age at death. Except for higher amounts of α-syn aggregates found in PD pericytes, the clearance kinetics are mainly similar indicating that pericytes in culture can retain certain characteristics specific to the brain from which they were isolated. This raises the possibility of an adaptive response in the PD pericytes where the previous priming to α-syn aggregates increases subsequent uptake upon repeated exposure. Furthermore, in PD, the gradual loss of pericytes is not uniform but affects every capillary differently. As such, individual pericytes derived from PD can have slightly different characteristics influencing degradation, which could explain the higher variability seen within PD pericytes [15].

Unique to this study is the prolonged timeframe (21 days post-exposure) that pericytes were followed. This prolonged follow-up period showed that cleavage occurs early after exposure, but that cleaved α-syn fragments remain present within pericytes for several weeks. It is important to highlight that the proportion of cells (17-59%) containing primarily cleaved α-syn at the later timepoints (14-21 days) can be relatively high depending on the strain. α-syn cleavage is positively correlated with accelerated aggregation and pathology in cell and mouse models [36–38]. Therefore, the spread of the remnant α-syn fragments through tunnelling nanotubes [22,39] or endosomal vesicles to neighbouring cells as seen in microglia [15,41] might still be possible.

Although these mechanisms are believed to dilute the burden over multiple cells[15], when exposure exceeds breakdown capacity, they may also contribute to prolonged presence of incompletely processed α-syn aggregates that possess seeding propensity given they retain their amyloid core. Indeed, all structural studies have demonstrated the C-terminal end of α-syn spanning residues 100 to 140 is moistly dynamic with little contribution to the fibrils amyloid core.

## CONCLUSION

Our data suggests that C-terminally processed α-syn strains remain present for prolonged periods adding to the current literature that insufficient degradation caused by high aggregated α-syn load could potentially favour propagation. Furthermore, pericytes could play an integral part in degrading α-syn aggregates within the brain and that this response is modified in PD. As such pericytes are a potential new target to reduce α-syn induced toxicity, slow down spread and delay disease onset.

## Acknowledgements

We would like to acknowledge the generosity of the brain donors and their family for their generous gift of brain tissue for research. We also thank Marika Eszes at the Neurological Foundation Human Brain Bank and all technical staff involved in the collection and processing of the human brain tissue at the Centre for Brain Research. We also thank Sheryl Feng and all staff of the Hugh Green Biobank for establishing the pericyte lines. This work was supported by the Hugh Green Foundation, Ian and Sue Parton. BVD was funded by the Michael J Fox Foundation (16420), the NeuroResearch Charitable Trust and Health Research Council Hercus (21/034). BH is funded by Brain Research New Zealand (grant number: 3715777). TP was supported by the Neurological Foundation grant (grant number: 1833PG) and the Douglas Charitable Trust. RM lab is supported by France Parkinson and the European Union Joint Programme on Neurodegenerative Disease Research and Agence National de la Recherche (contracts PROTEST-70, ANR-17-JPND-0005-01 and Trans-PathND, ANR-17-JPND-0002-02). MD is supported through a Programme grant from the Health Research Council of New Zealand (21/730) and the Michael J Fox Foundation (16420).

## Author Contributions

BVD, RM, BH, TS and MD contributed to the conception and design of the experiments. Experiments were performed by BVD, TS, VL. Preparation quality control of α-syn aggregates was done by AA, TB, RM. Primary human pericytes were provided by MD. Human tissue was provided by MAC, RLM. Primary human brain cells were provided by TP, JC, PS, MD. Data collection and analysis were performed by BVD and BH. The manuscript was written by BVD, BH, RM. All authors read and approved the final manuscript.

## Supplementary material

**Figure S 1.**
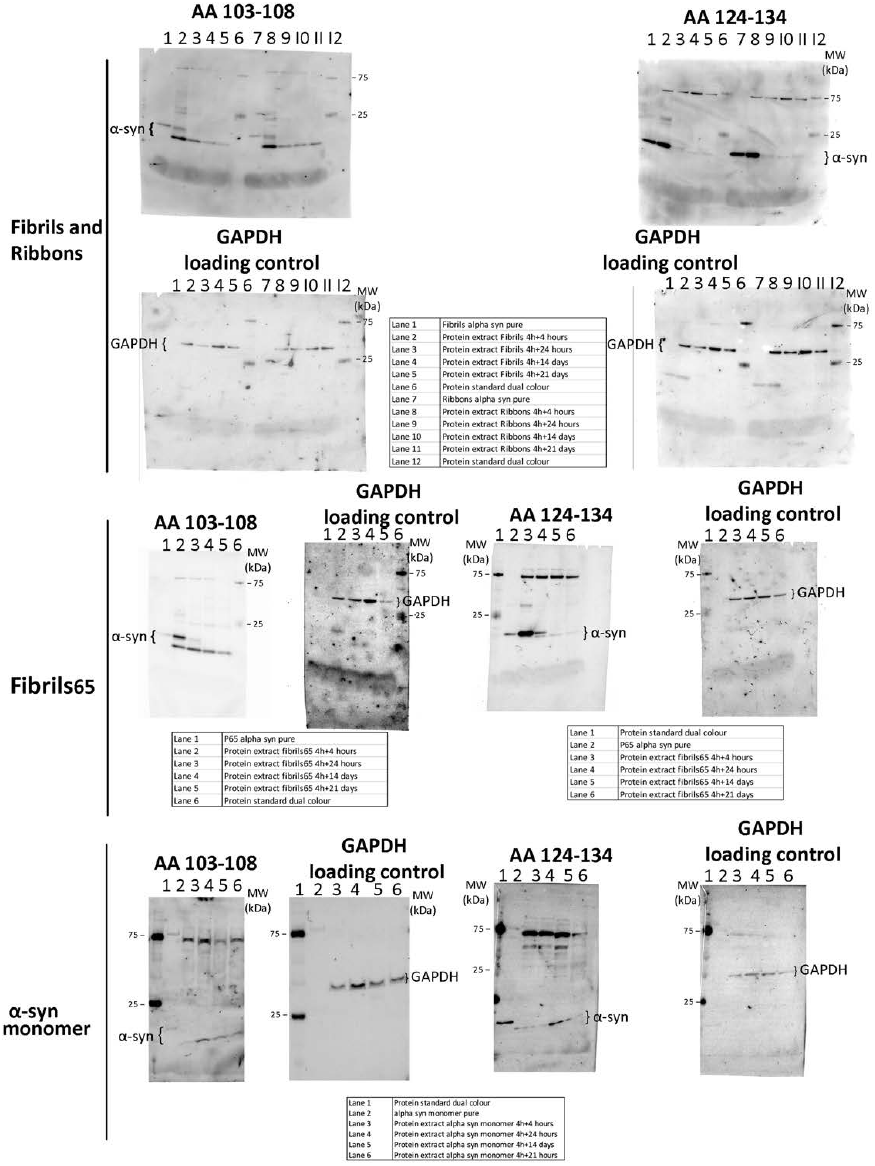
Full western blots for Fibrils, Ribbons and fibrils65 with AA103-108, AA124-134 and GAPDH. Two blots used for each alpha synuclein strain. Each blot was labelled with α-syn antibodies (AA103-108 or AA124-134), imaged, stripped and relabelled for GAPDH.

**Figure S 2.**
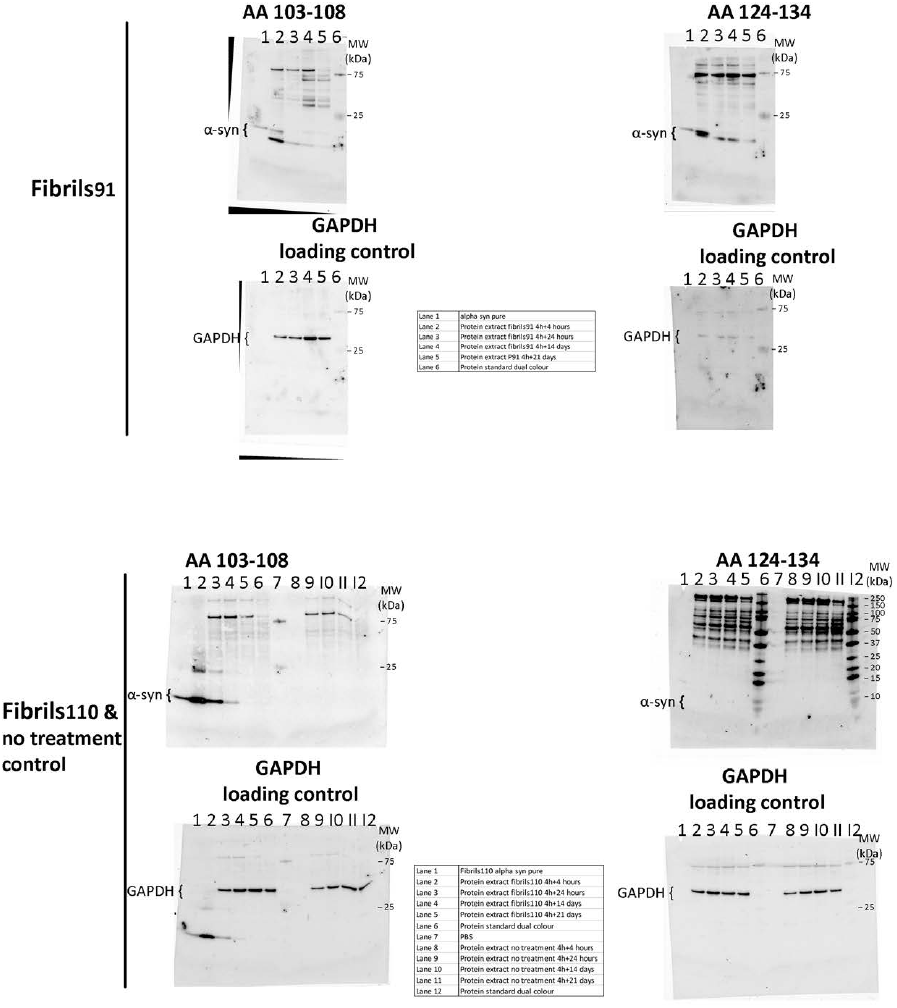
Full western blots for fibrils91, fibrils110 and no treatment control with AA103-108, AA124-134 and GAPDH. Two blots used for each alpha synuclein strain. Each blot was double labelled with α-syn antibodies (AA103-108 or AA124-134), imaged, stripped and relabelled for GAPDH.

**Figure S 3.**
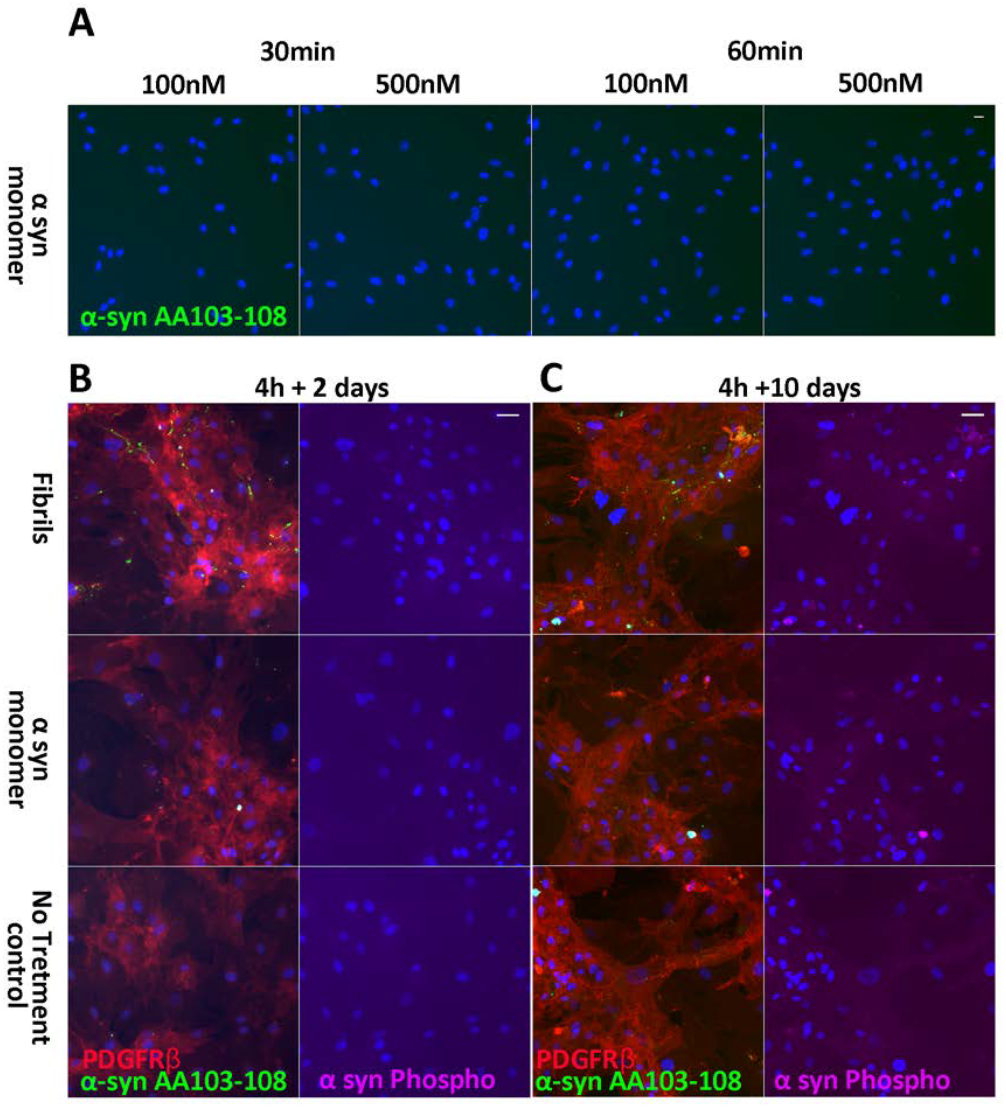
Immunofluorescent labelling of endogenous, monomeric or fibrillar Fibrils treated pericytes. No α-syn puncta detected with α-syn specific antibodies AA103-108 (green) after treatment with monomeric α-syn (100µM and 500µM) (A). Immunofluorescent labelling of endogenous, monomeric or fibrillar Fibrils treated pericytes with α-syn specific antibodies AA103-108 (green), PDGFRβ (red) and α-syn Phospho S129 (magenta) after 4 hour pre-treatment with 100nM Fibrils or monomeric α-syn (2 days, B; 10 days C). Scale bars represent 20 µm.

**Figure S 4.**
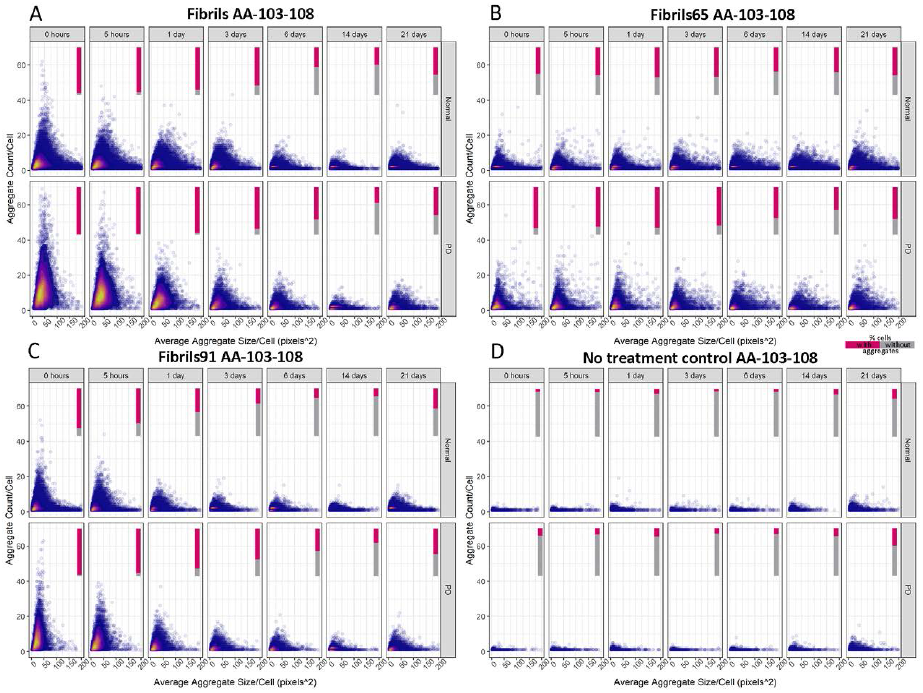
Density plots of showing a differential response to α-syn aggregates between control and PD-derived pericytes based on AA103-108 for Fibrils, FIBRILS65, FIBRILS91 no treatment control. Cells without aggregates are excluded from density plots. Relative amounts of cells with aggregates represented in bar (% cells with aggregates in magenta, % cells without aggregates in grey). The colours represent a 2D kernel density estimation. It is scaled to 1 for each graph, with the bright yellow area displaying the highest density of cells.

Movie S 1 Life cell imaging (120 hours) of pericytes pre-treated for 4 hours with 100nM Fibrils-atto647.

Movie S 2 Life cell imaging (120 hours) of pericytes pre-treated for 4 hours with 100nM Ribbons-atto647.

Movie S 3 Life cell imaging (120 hours) of pericytes pre-treated for 4 hours with 100nM fibrils65-atto647.

Movie S 4 Life cell imaging (72 hours) of pericytes pre-treated for 4 hours with 100nM fibrils91-atto647.

Movie S 5 Life cell imaging (90 hours) of pericytes pre-treated for 4 hours with 100nM fibrils110-atto647.

Movie S 6 Life cell imaging (90 hours) of mixed neuronal cell cultures pre-treated for 4 hours with 100nM fibrils110-atto647

Movie S 7 Life cell imaging showing α-syn uptake by pericyte adjacent to microglia

Supplementary M&M 1 R script used for Image analysis

## References

1 Villar-Piqué A, Lopes da Fonseca T, Outeiro TF. Structure, function and toxicity of alpha-synuclein: the Bermuda triangle in synucleinopathies. Journal of Neurochemistry. 2016;139:240–55.

2 Malmberg M, Malm T, Gustafsson O, Sturchio A, Graff C, Espay AJ, et al. Disentangling the Amyloid Pathways: A Mechanistic Approach to Etiology. Frontiers in Neuroscience. 2020;14(April):1–11.

3 Stefanis L, Emmanouilidou E, Pantazopoulou M, Kirik D, Vekrellis K, Tofaris GK. How is alpha-synuclein cleared from the cell? Journal of Neurochemistry. 2019;150(5):577–90.

4 Braak H, Tredici K del, Rüb U, de Vos RAI, Jansen Steur ENH, Braak E. Staging of brain pathology related to sporadic Parkinson’s disease. Neurobiol Aging. 2003;24(2):197–211.

5 Tran J, Anastacio H, Bardy C. Genetic predispositions of Parkinson’s disease revealed in patient-derived brain cells. npj Parkinson’s Disease. 2020;6(1). DOI: 10.1038/s41531-020-0110-8

6 Hoppe SO, Uzunoğlu G, Nussbaum-Krammer C. α-Synuclein Strains: Does Amyloid Conformation Explain the Heterogeneity of Synucleinopathies? Biomolecules. 2021;11(7). DOI: 10.3390/biom11070931

7 Van der Perren A, Gelders G, Fenyi A, Bousset L, Brito F, Peelaerts W, et al. The structural differences between patient-derived α-synuclein strains dictate characteristics of Parkinson’s disease, multiple system atrophy and dementia with Lewy bodies. Acta Neuropathologica. 2020;139(6):977–1000.

8 Peelaerts W, Bousset L, van der Perren A, Moskalyuk A, Pulizzi R, Giugliano M, et al. α-Synuclein strains cause distinct synucleinopathies after local and systemic administration. Nature. 2015;522(7556):340–4.

9 Lau A, So RWL, Lau HHC, Sang JC, Ruiz-Riquelme A, Fleck SC, et al. α-Synuclein strains target distinct brain regions and cell types. Nature Neuroscience. 2020;23(1):21–31.

10 Liu D, Guo J, Su J, Svanbergsson A, Yuan L, Haikal C, et al. Differential seeding and propagating efficiency of α-synuclein strains generated in different conditions. Translational Neurodegeneration. 2021;10(20):1–15.

11 Sorrentino ZA, Giasson BI, Chakrabarty P. α-Synuclein and astrocytes: tracing the pathways from homeostasis to neurodegeneration in Lewy body disease. Acta Neuropathologica. 2019;138(1):1–21.

12 Tanriöver G, Bacioglu M, Schweighauser M, Mahler J, Wegenast-Braun BM, Skodras A, et al. Prominent microglial inclusions in transgenic mouse models of α-synucleinopathy that are distinct from neuronal lesions. Acta Neuropathologica Communications. 2020;8(1):1–11.

13 Stevenson TJ, Murray HC, Turner C, Faull RLM, Dieriks B v, Curtis MA. α-synuclein inclusions are abundant in non-neuronal cells in the anterior olfactory nucleus of the Parkinson’ s disease olfactory bulb. Scientific Reports. 2020;1–10.

14 Loria F, Vargas JY, Bousset L, Syan S, Salles A, Melki R, et al. α-Synuclein transfer between neurons and astrocytes indicates that astrocytes play a role in degradation rather than in spreading. Acta Neuropathologica. 2017 Jul;1–20.

15 Scheiblich H, Dansokho C, Mercan D, Schmidt S v., Bousset L, Wischhof L, et al. Microglia jointly degrade fibrillar alpha-synuclein cargo by distribution through tunneling nanotubes. Cell. 2021;184(20):5089-5106.e21.

16 Russ K, Teku G, Bousset L, Redeker V, Piel S, Savchenko E, et al. TNF-α and α-synuclein fibrils differently regulate human astrocyte immune reactivity and impair mitochondrial respiration. Cell Reports. 2021;34(12):108895.

17 Rustenhoven J, Jansson D, Smyth LC, Dragunow M. Brain Pericytes As Mediators of Neuroinflammation. Trends in Pharmacological Sciences. 2017;38(3):291–304.

18 Jansson D, Rustenhoven J, Feng S, Hurley D, Oldfield RL, Bergin PS, et al. A role for human brain pericytes in neuroinflammation. Journal of Neuroinflammation. 2014;11:104.

19 Rustenhoven J, Scotter EL, Jansson D, Kho DT, Oldfield RL, Bergin PS, et al. An anti-inflammatory role for C/EBPδ in human brain pericytes. Scientific Reports. 2015;5:12132.

20 Gray MT, Woulfe JM. Striatal blood-brain barrier permeability in Parkinson’s disease. Journal of Cerebral Blood Flow and Metabolism. 2015;35(5):747–50.

21 Yang P, Waldvogel HJ, Faull RLM, Dragunow M, Guan J. Vascular degeneration in Parkinson ’ s disease. Alzheimer’s and Parkinson’s Disease. 2016;2(1).

22 Dieriks BV, Park TI-H, Fourie C, Faull RLM, Dragunow M, Curtis MA. α-synuclein transfer through tunneling nanotubes occurs in SH-SY5Y cells and primary brain pericytes from Parkinson’s disease patients. Scientific Reports. 2017 Feb;7:42984.

23 Ghee M, Melki R, Michot N, Mallet J. PA700, the regulatory complex of the 26S proteasome, interferes with α-synuclein assembly. FEBS J. 2005;272(16):4023–33.

24 Bousset L, Pieri L, Ruiz-Arlandis G, Gath J, Jensen PH, Habenstein B, et al. Structural and functional characterization of two alpha-synuclein strains. Nature Communications. 2013;4. DOI: 10.1038/ncomms3575

25 Makky A, Bousset L, Polesel-Maris J, Melki R. Nanomechanical properties of distinct fibrillar polymorphs of the protein α-synuclein. Scientific Reports. 2016;6:1–10.

26 Pieri L, Madiona K, Melki R. Structural and functional properties of prefibrillar α-synuclein oligomers. Scientific Reports. 2016;6(April):1–15.

27 Grozdanov V, Bousset L, Hoffmeister M, Bliederhaeuser C, Meier C, Madiona K, et al. Increased Immune Activation by Pathologic α-Synuclein in Parkinson’s Disease. Annals of Neurology. 2019;86(4):593–606.

28 Park TI-H, Smyth LCD, Aalderink M, Woolf ZR, Rustenhoven J, Lee K, et al. Routine culture and study of adult human brain cells from neurosurgical specimens. Nat Protoc. 2022;1–32.

29 Rustenhoven J, Smyth LC, Jansson D, Schweder P, Aalderink M, Scotter EL, et al. Modelling physiological and pathological conditions to study pericyte biology in brain function and dysfunction. BMC Neuroscience. 2018;19(1):1–15.

30 Rustenhoven J, Aalderink M, Scotter EL, Oldfield RL, Bergin PS, Mee EW, et al. TGF-beta1 regulates human brain pericyte inflammatory processes involved in neurovasculature function. Journal of Neuroinflammation. 2016;13(1):1–15.

31 Smyth LCD, Rustenhoven J, Park TI-H, Schweder P, Jansson D, Heppner PA, et al. Unique and shared inflammatory profiles of human brain endothelia and pericytes. Journal of Neuroinflammation. 2018;15(1):138.

32 Smyth LCD, Rustenhoven J, Scotter EL, Schweder P, Faull RLM, Park TIH, et al. Markers for human brain pericytes and smooth muscle cells. Journal of Chemical Neuroanatomy. 2018;92(January):48– 60.

33 Park TI-H, Schweder P, Lee K, Dieriks B V, Jung Y, Smyth L, et al. Isolation and culture of functional adult human neurons from neurosurgical brain specimens. Brain Communications. 2020;2(2). DOI: 10.1093/braincomms/fcaa171

34 Waldvogel HJ, Curtis MA, Baer K, Rees MI, Faull RL. Immunohistochemical staining of post-mortem adult human brain sections. Nat Protoc. 2006;1(6):2719–32.

35 Highet B, Dieriks BV, Murray HC, Faull RLM, Curtis MA. Huntingtin aggregates in the olfactory bulb in Huntington’s disease. Frontiers in Aging Neuroscience. 2020;12:261.

36 Shrivastava AN, Bousset L, Renner M, Redeker V, Savistchenko J, Triller A, et al. Differential Membrane Binding and Seeding of Distinct α-Synuclein Fibrillar Polymorphs. Biophysical Journal. 2020;118(6):1301–20.

37 Duan L, Zhang X-D, Miao W-Y, Sun Y-J, Xiong G, Wu Q, et al. PDGFRβ Cells Rapidly Relay Inflammatory Signal from the Circulatory System to Neurons via Chemokine CCL2. Neuron. 2018;100(1):183-200.e8.

38 Shahmoradian SH, Lewis AJ, Genoud C, Hench J, Moors TE, Navarro PP, et al. Lewy pathology in Parkinson’s disease consists of crowded organelles and lipid membranes. Nature Neuroscience. 2019;22(7):1099–109.

39 Lee HJ, Suk JE, Bae EJ, Lee SJ. Clearance and deposition of extracellular α-synuclein aggregates in microglia. Biochemical and Biophysical Research Communications. 2008;372(3):423–8.

40 Grozdanov V, Danzer KM. Release and uptake of pathologic alpha-synuclein. Cell and Tissue Research. 2018;373(1):175–82.

41 Xia Y, Zhang G, Kou L, Yin S, Han C, Hu J, et al. Reactive microglia enhance the transmission of exosomal α-synuclein via toll-like receptor 2. Brain. 2021;144(7):2024–37.

42 Pieri L, Chafey P, le Gall M, Clary G, Melki R, Redeker V. Cellular response of human neuroblastoma cells to α-synuclein fibrils, the main constituent of Lewy bodies. Biochimica et Biophysica Acta (BBA)-General Subjects. 2016;1860(1):8–19.

43 Moors TE, Maat CA, Niedieker D, Mona D, Petersen D, Timmermans-Huisman E, et al. Subcellular orchestration of alpha-synuclein variants in Parkinson’s disease brains revealed by 3D multicolor STED microscopy. bioRxiv. 2018 DOI: 10.1101/470476

44 Sorrentino ZA, Giasson BI. The emerging role of α-synuclein truncation in aggregation and disease. Journal of Biological Chemistry. 2020;295(30):10224–44.

45 Bell RD, Winkler EA, Sagare AP, Singh I, LaRue B, Deane R, et al. Pericytes Control Key Neurovascular Functions and Neuronal Phenotype in the Adult Brain and during Brain Aging. Neuron. 2010 DOI: 10.1016/j.neuron.2010.09.043

46 Kristensson K, Olsson Y. Accumulation of protein tracers in pericytes of the central nervous system following systemic injection in immature mice. Acta Neurologica Scandinavica. 1973 Jan;49(2):189– 94.

47 Schultz N, Byman E, Fex M, Wennström M. Amylin alters human brain pericyte viability and NG2 expression. Journal of Cerebral Blood Flow and Metabolism. 2017 DOI: 10.1177/0271678X16657093

48 Cuervo AM. Impaired Degradation of Mutant -Synuclein by Chaperone-Mediated Autophagy. Science (1979). 2004 Aug;305(5688):1292–5.

49 Emmanouilidou E, Stefanis L, Vekrellis K. Cell-produced α-synuclein oligomers are targeted to, and impair, the 26S proteasome. Neurobiology of Aging. 2010 DOI: 10.1016/j.neurobiolaging.2008.07.008

50 Dargemont C, Ossareh-Nazari B. Cdc48/p97, a key actor in the interplay between autophagy and ubiquitin/proteasome catabolic pathways. Biochimica et Biophysica Acta - Molecular Cell Research. 2012;1823(1):138–44.

51 Xilouri M, Brekk OR, Stefanis L. α-Synuclein and protein degradation systems: a reciprocal relationship. Mol Neurobiol. 2013 DOI: 10.1007/s12035-012-8341-2

52 Yang P, Waldvogel H, Turner C, Faull R, Dragunow M, Guan J. Vascular Remodelling is Impaired in Parkinson Disease. Journal of Alzheimer’s and Parkinsonism. 2017;7(2). DOI: 10.4172/2161-0460.1000313

53 Daher JPL, Ying M, Banerjee R, McDonald RS, Hahn MD, Yang L, et al. Conditional transgenic mice expressing C-terminally truncated human α-synuclein (αSyn119) exhibit reduced striatal dopamine without loss of nigrostriatal pathway dopaminergic neurons. Molecular Neurodegeneration. 2009;4(1). DOI: 10.1186/1750-1326-4-34

54 Tofaris GK, Reitböck PG, Humby T, Lambourne SL, O’Connell M, Ghetti B, et al. Pathological changes in dopaminergic nerve cells of the substantia nigra and olfactory bulb in mice transgenic for truncated human α-synuclein(1-120): Implications for lewy body disorders. Journal of Neuroscience. 2006;26(15):3942–50.

55 Chakroun T, Evsyukov V, Nykänen NP, Höllerhage M, Schmidt A, Kamp F, et al. Alpha-synuclein fragments trigger distinct aggregation pathways. Cell Death and Disease. 2020;11(2). DOI: 10.1038/s41419-020-2285-7

56 Alarcon-Martinez L, Villafranca-Baughman D, Quintero H, Kacerovsky JB, Dotigny F, Murai KK, et al. Interpericyte tunnelling nanotubes regulate neurovascular coupling. Nature. 2020;585(7823):91–5.

